# The photobiology of the human circadian clock

**DOI:** 10.1101/2021.10.13.463655

**Authors:** Robin A. Schoonderwoerd, Mischa de Rover, Jan A.M. Janse, Lydiane Hirschler, Channa R. Willemse, Leonie Scholten, Ilse Klop, Sander van Berloo, Matthias J.P. van Osch, Dick F. Swaab, Johanna H. Meijer

**Author notes:** = Corresponding author.

## Abstract

In modern society, the widespread use of artificial light at night disrupts the suprachiasmatic nucleus (SCN), which serves as our central circadian clock. Existing models describe excitatory responses of the SCN to primarily blue light, but direct measures in humans are absent. The combination of state-of-the-art neuroimaging techniques and custom-made MRI compatible LED devices allowed to directly measure the light response of the SCN. In contrast to the general expectation, we found that SCN activity was suppressed by light. The suppressions were observed not only in response to narrowband blue light (λ_max_: 470nm) but remarkably, also in response to green (λ_max_: 515nm) and orange (λ_max_: 590nm), but not to violet light (λ_max_: 405nm). The broadband sensitivity of the SCN implies that strategies on light exposure should be revised: enhancement of light levels during daytime is possible with wavelengths other than blue, while during nighttime, all colors are potentially disruptive.

## Introduction

Due to the Earth’s rotation around its axis, many organisms developed an internal clock to anticipate the predictable changes in the environment that occur every 24 hours, including the daily light-dark cycle. In mammals, this clock is located in the suprachiasmatic nucleus (SCN), located in the hypothalamus directly above the optic chiasm.^1,2^ The SCN receives information from the retina regarding ambient light levels via intrinsically photosensitive retinal ganglion cells (ipRGCs), thus synchronizing its internal clock to the external light-dark cycle. ipRGCs contain the photopigment melanopsin, which is maximally sensitive to blue light, with a peak response to 480-nm light.^3,4^ In addition, ipRGCs also receive input from rod cells and cone cells.^5–7^ The three cone cell subtypes in the human retina respond maximally to 420-nm, 534-nm, and 563-nm light, while rod cells respond maximally to 498-nm light.^8^ In rodents, input from cone cells renders the SCN sensitive to a broad spectrum of wavelengths,^9^ while rod cells mediate the SCN’s sensitivity to low-intensity light.^10,11^ Recently, these findings in rodents were proposed to translate to humans,^12^ suggesting that the human clock is not only sensitive to blue light, but may also be sensitive to other colors.

In humans, circadian responses to light are generally measured indirectly, for example by measuring melatonin levels or 24-hour behavioral rhythms. These indirect measures revealed that circadian responses to light in humans are most sensitive to blue light;^13–16^ however, green light has also been found to contribute to circadian phase shifting and changes in melatonin to a larger extent than would have been predicted based solely on the melanopsin response, suggesting that rods and/or cones may also provide functional input to the circadian system in humans.^17^ Despite this indirect evidence suggesting that several colors can affect the human circadian clock, this has never been measured directly due to technical limitations. Thus, current guidelines regarding the use of artificial light are based solely on the clock’s sensitivity to blue light. For example, blue light is usually filtered out in electronic screens during the night,^18,19^ and blue-enriched light is used by night shift workers to optimize their body rhythm for achieving maximum performance.^20–22^

The ability to directly image the human SCN *in vivo* has been severely limited due to its small size and the relatively low spatial resolution provided by medical imaging devices. Previous functional magnetic resonance imaging (fMRI) studies using 3-Tesla (3T) scanners were restricted to recording the “suprachiasmatic area”, which encompasses a large part of the hypothalamus and thus includes many other potentially light-sensitive nuclei.^23–25^ To overcome this limitation, we used a 7T MRI scanner, which can provide images with sufficiently high spatial resolution to image small brain nuclei^26^ such as the SCN. Here, we applied colored light stimuli to healthy volunteers using a custom-designed MRI-compatible light-emitting diode (LED) device designed to stimulate specific photoreceptors while measuring SCN activity using fMRI. Using new analytical approaches, we then identified the SCN’s response, the smallest brain nucleus that has so far been imaged. We found that the human SCN responds to a broad range of wavelengths, i.e. blue, green and orange light. Surprisingly, we also found that SCN activity is actually suppressed—not activated—by light.

## Results

First, we developed an MRI-compatible LED device designed to apply specific wavelengths of light to the participants’ eyes while measuring brain activity using fMRI (Figure 1A, B). The LED device was attached to the inside of the head coil, providing sufficiently high light intensity and whole-field retinal illumination, which is important given that melanopsin-expressing cells are present throughout the retina.^27^ Moreover, the LED device was designed using non-ferromagnetic materials and is therefore unaffected by high-frequency radio pulses and the strong magnetic field produced by the MRI scanner (see Methods).

**Figure 1:**
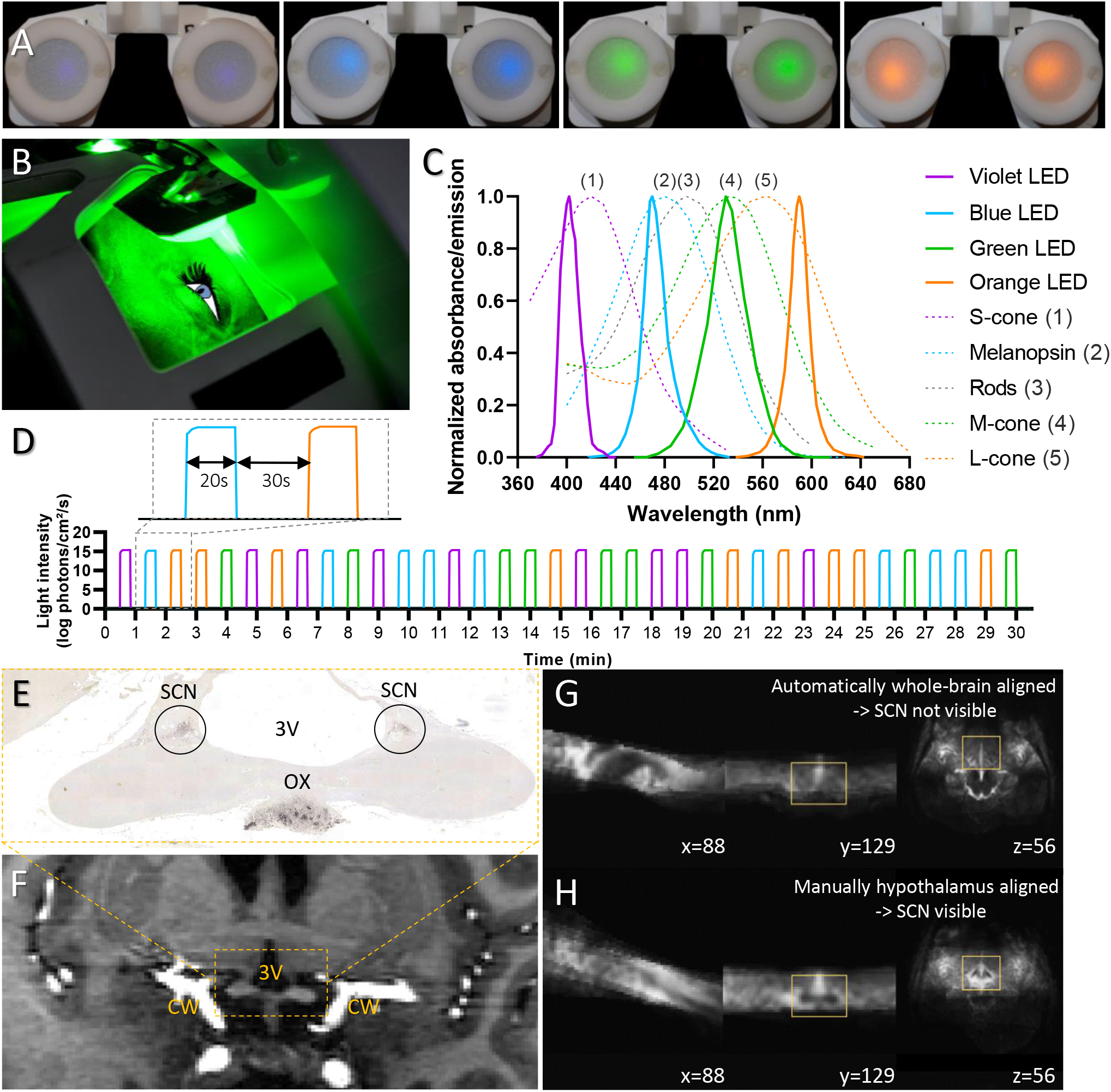
The fMRI-compatible LED device, light protocol, and SCN localization. (A, B) A set of MRI-compatible light-emitting glasses. (C) LEDs were used to stimulate the indicated human photoreceptors close to their known peak absorption spectra; L-cone, M-cone, and S-cone refer to long-, medium-, and short-wavelength cone cells, respectively. (D) Light protocol: 20-sec light pulses followed by 30-sec periods of darkness. (E) Vasopressin-stained sections from human post-mortem brains. OX, optic chiasm; 3V, third ventricle. (F) SCN is indicated by the dashed yellow rectangle in a high-resolution T1-weighted image. CW, circle of Willis. (G, H) Improvement of chiasm alignment (within yellow rectangles) following manual registration.

To evaluate the output performance of the LED device, we first measured the individual emission spectra of the four sets of LEDs. Each set of LEDs emitted light at the expected wavelength, with minimum overlap with the absorption spectra of other receptors (Figure 1C). Next, we configured the protocol for the LED device to deliver 15.2 log photons/cm^2^/s via each individual LED set using the timing shown in Figure 1D (see Methods for details).

For functional imaging, the location of both SCN was determined by comparing each participant’s anatomical images with reference images based on immunocytochemical staining of vasopressin in sections prepared from human post-mortem brains (Figure 1E). Because the SCN are located in close proximity to both the large blood vessels of the circle of Willis and the third ventricle (Figure 1E, F), we used the Elastix registration toolbox (https://elastix.lumc.nl/) to correct the functional data for motion, and we used ICA-AROMA (Independent Component Analysis - Automatic Removal of Motion Artifacts) for de-noising of any breathing and heart rate–associated artifacts. To register the functional data slices to the standard MNI152 template, automatic registration focusing on the whole-brain alignment was insufficient to visualize the SCN and optic chiasm in the mean plot (Figure 1G); therefore, we performed manual registration focusing on the SCN region and only in the manually aligned averages of the functional data, the outline of the optic chiasm emerged in the mean plot (Figure 1H).

For our study, all participants were male, as post-mortem studies indicate that the SCN in men is spherical, compared to the more elongated SCN in women,^28^ thus increasing our ability to capture the SCN signal, given the shape and size of the voxels. All 12 participants were healthy male adults (30 ± 12 years of age; range: 22-58 years), with an MSFsc (mid-sleep time on free days, corrected for sleep debt) value of 04:32 ± 01:30 relative to midnight, thus there was no overrepresentation of extreme morning or evening types. All scans were performed between 22:00 (10:00 pm) and 23:00 (11:00 pm), as previous studies suggest that light elicits the strongest SCN response at night.^23,29^

To characterize the light-induced response in the SCN, MRI masks were drawn *a priori* covering both the left and right SCN based exclusively on anatomical localization (Figure 2A) by comparing the anatomical scans with the immunocytochemical staining shown in Figure 1E. Time series data were extracted, and the blood oxygen level-dependent (BOLD) signal was averaged for light stimuli when all four colors were combined (i.e., the responses elicited by all 36 light pulses were averaged). As a positive control, masks of equal size were also drawn in the visual cortex V1 region (Figure 2B), and time series data were extracted to generate the average waveform (Figure 2E). As a negative control, masks of the same size were also drawn in a region containing cerebrospinal fluid (CSF) (Figure 2C, F and Supplemental Figure S2). All twelve participants were included in the SCN and CSF analyses, whereas only nine participants were included in the V1 analysis, as the functional scan did not cover the masked areas of V1 in three participants. The light-induced responses were quantified by averaging the percent change in the BOLD signal for the last 15 seconds of each 20-sec light pulse and are summarized in Figure 2G. We found that on average, all 36 light pulses significantly reduced the BOLD signal in the SCN by 0.38±0.10 percent (*p*=0.003) and significantly increased the BOLD signal in the V1 region of the visual cortex by 1.45±0.27 percent (*p*<0.001); as expected, the light pulses had no effect on the BOLD signal measured in the CSF (*p*=0.84). A mixed-effects model showed significant differences between the three regions (F(1.443, 13.71) = 39.24; *p*<0.0001), and Šidák’s multiple comparisons test showed a significant difference in the responses measured between the SCN and CSF (*p*=0.039).

**Figure 2:**
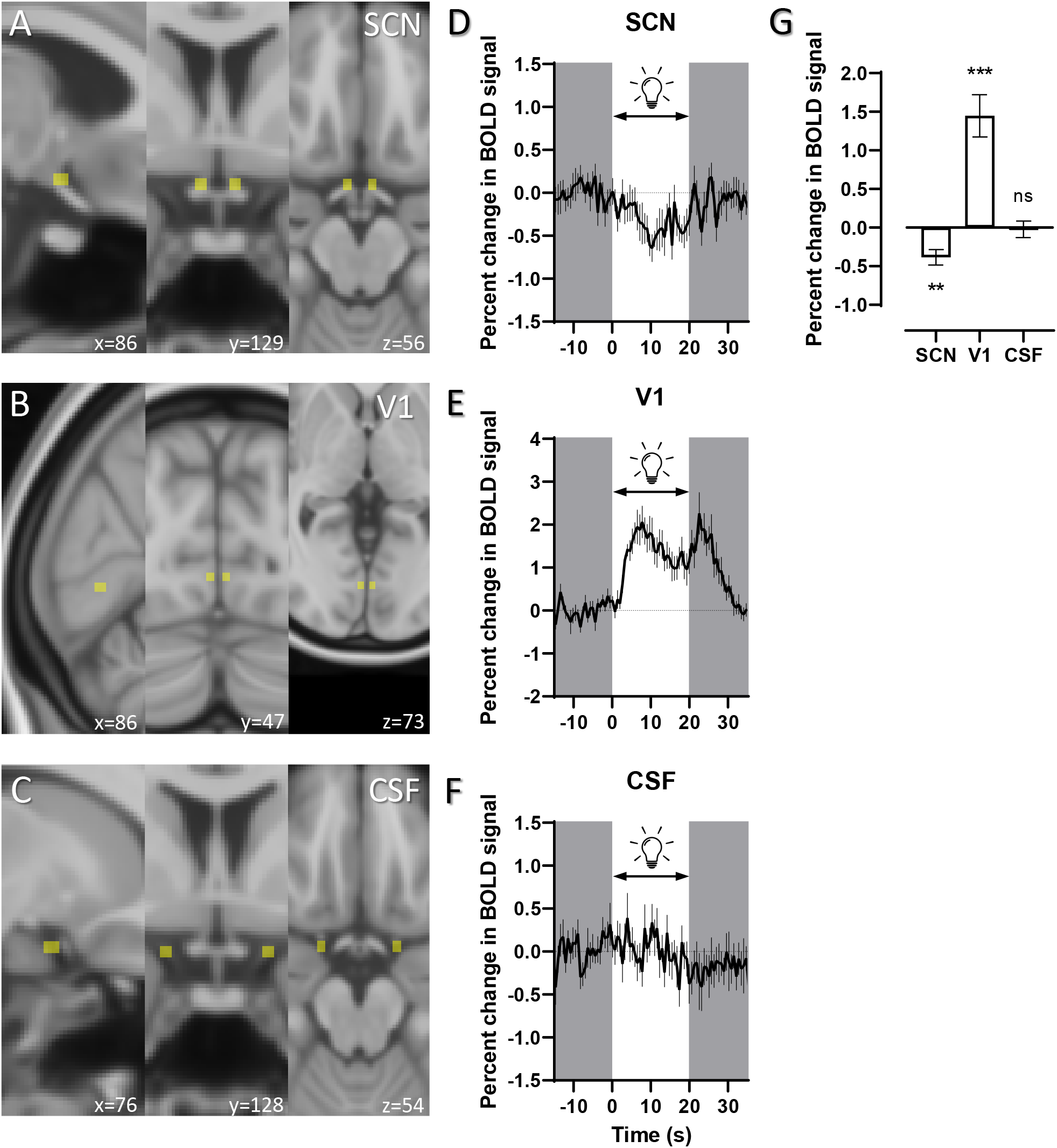
Light causes a suppression in BOLD signal in the SCN. (A) Masks (3×4×3 mm) were drawn on the standard MNI152 template (yellow shaded squares) for the SCN, (B) V1 and (C) CSF. (D-F) Average time course of the change in BOLD signal measured in the SCN (D), visual cortex region V1 (E), and CSF (F) in all 12 participants. Darkness was indicated by the gray background. (G) Summary of the average change in the BOLD signal measured in the SCN, VI region, and CSF in response to all 36 light pulses. \*\**p*<0.01, \*\*\**p*<0.001, and ns, not significant (one-sample *t*-test). In this and subsequent figures, all summary data are presented as the mean ± SEM.

We then used a general linear model (GLM) to perform a voxel-based analysis of the responses to individual colors and found voxels showing a significant response (*p*<0.01, uncorrected) at the location of the SCN, bilaterally above the optic chiasm (Fig. 3). Interestingly, although violet light did not cause a significant change in the BOLD signal in the SCN (Figure 3A, E) blue light (Figure 3B, F), green light (Figure C, G), and orange light (Figure D, H) each induced a significant suppression in the BOLD signal (Figure 3I). All voxels at the SCN location had a z-stat value of <2.3, indicating that these light-induced responses in the SCN reflect a suppression in neuronal activity.

**Figure 3:**
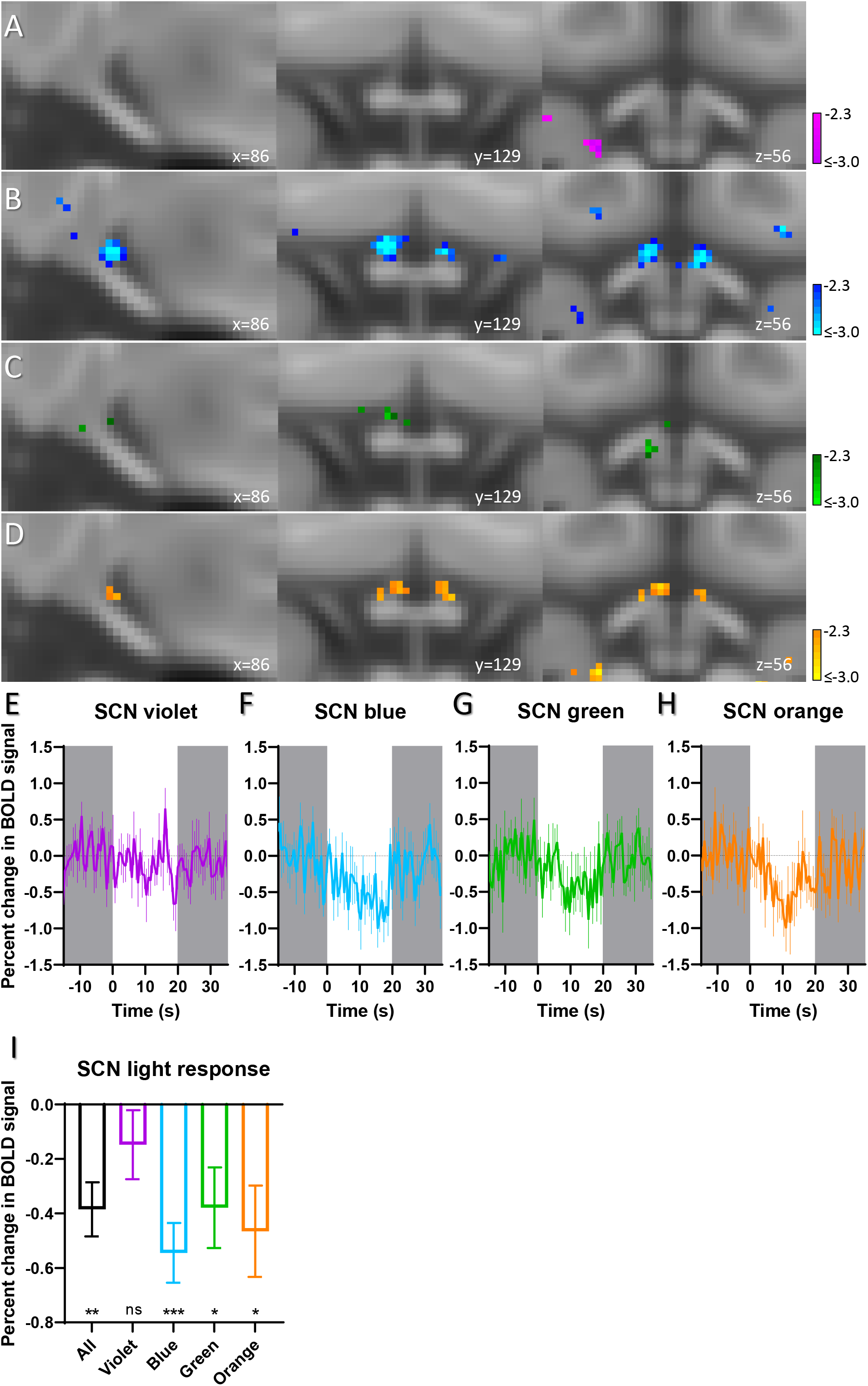
The SCN responds to several wavelengths in addition to blue light. (A-D) Light-responsive voxels showing the BOLD signal measured in the SCN of all 12 participants in response to violet (A), blue (B), green (C), and orange (D) light (*p*<0.01 uncorrected for illustration). (E-H) Average time course of the change in BOLD signal measured in the SCN. Percent change in BOLD signal following violet (E), blue (F), green (G), and orange (H) light. (I) Average light responses and significances. Data in the first bar (in black) were reproduced from Figure 2G for comparison purposes. \**p*<0.05, \*\**p*<0.01, \*\*\**p*<0.001, and ns, not significant (one-sample *t*-test). A one-way repeated measures ANOVA revealed no significant differences in the response between colors (F(2,764, 30,41) = 2,292; *p*=0.102).

We then performed the same analysis in the V1 region of the visual cortex as a positive control (n=9 participants) (Figure 4). As expected, all four individual colors induced a significant increase (*p*<0.01, uncorrected) in the BOLD signal (Figure 4I).

**Figure 4:**
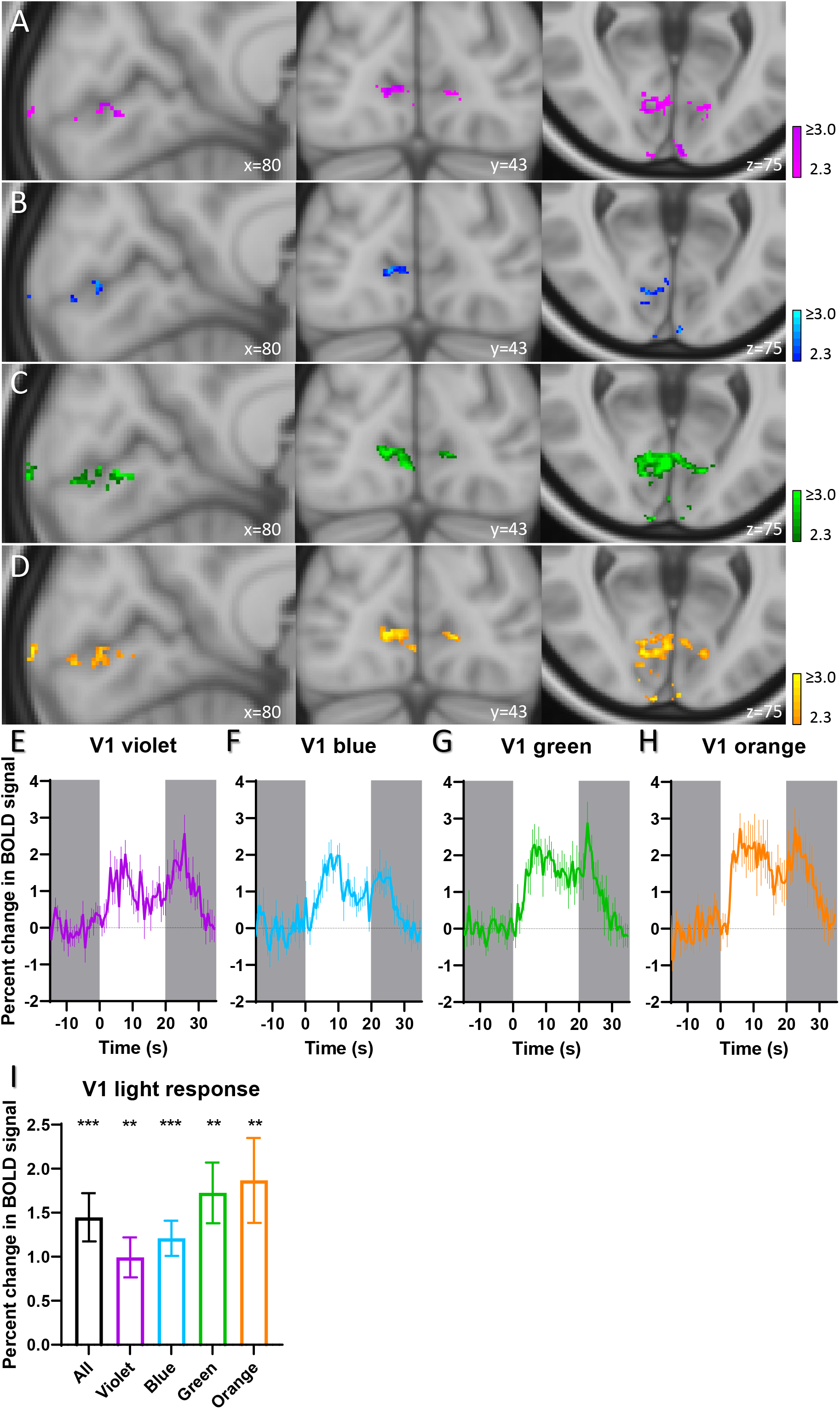
All four colors induce a response in the visual cortex. (A-D) Light-responsive voxels showing the BOLD signal measured in the V1 region of 9 participants in response to violet (A), blue (B), green (C), and orange (D) light (*p*<0.01 uncorrected for illustration). (E-H) Average time course of the change in BOLD signal measured in the V1 region. Percent change in BOLD signal following violet (E), blue (F), green (G), and orange (H) light. (I) Summary of the average change in the BOLD signal measured in the V1 region. The data in the first bar were reproduced from Figure 2G for comparison purposes. \*\**p*<0.01 and \*\*\**p*<0.001 (one-sample *t*-test). One-way repeated measures ANOVA shows no significant difference between colors (F(1.410, 11.28) = 3.665; *p*=0.071).

## Discussion

In humans, a properly functioning circadian clock is essential for maintaining health; however, our modern society can alter our circadian clock’s function, for example due to our excessive use of artificial light at night, long flights that span multiple time zones, and shift work (particularly rotating shift work). To preserve clock function, we should maximize our exposure to light during the day and minimize the harmful effects of artificial light during the night; thus, strategies designed to optimize our exposure to light—particularly light at the “correct” wavelength—are needed. As a major step toward developing such a strategy, we created an MRI-compatible LED device and directly measured activity in the participants’ SCN using state-of-the-art neuroimaging techniques while delivering specific colors of light. We found that the SCN responded negatively to blue light—and, surprisingly, to green and orange light as well— whereas violet light had no measurable effect. Given that violet light elicited the weakest effect in the visual cortex among all four colors, its lack of a significant effect in the SCN may reflect the partial filtering of short-wavelength light in the eye by the lens and the macular pigment.^30^

The differential contribution of specific photoreceptors to image-forming vision in the visual cortex versus non-image-forming vision in the SCN is reflected by the difference in responses to blue light between these two brain regions; specifically, we found that blue light elicited the strongest response in the SCN, yet elicited one of the weakest responses in the visual cortex. This relatively strong response to blue light in the SCN is likely due to melanopsin-mediated input.^4^ Furthermore, we found that green and orange light elicited similar responses in the SCN, providing strong evidence that the activity of L-cone cells in the human retina contributes to the SCN’s response to external light.

We found that blue, green, and orange light induced a negative change in the BOLD signal measured in the human SCN. These results are in contrast to a previous fMRI study by Vimal *et al*., in which the authors observed a positive change in the BOLD signal in response to light stimuli.^23^ It is important to note, however, that the authors analyzed a larger volume covering more than just the SCN, with lower spatial resolution (3.44×3.44×1.9 mm, compared to 1.25×1.25×1.65 mm in our study, i.e. more than a factor of 8 smaller voxel volume). More recently, McGlashan *et al*. measured the effect of light stimuli on the BOLD signal in the suprachiasmatic area and found that half of the measurements indicated a positive change in the BOLD signal, while the other half indicated a negative response; however, in their study the suprachiasmatic area encompassed the entire hypothalamus, which limited the interpretation of their results.^25^

The negative change in the BOLD signal in the SCN in response to light likely reflects an inhibitory effect of light input on neuronal activity,^31,32^ as recently reported for the habenula^33^ and amygdala^34^. Hypothalamic inhibition in the suprachiasmatic area in response to bright white light at night has been shown using positron emission tomography.^35^ The deactivated area in that study covered not only the SCN, but a large hypothalamic area, also including the subparaventricular zone and ventro-lateral preoptic area. In our study, the spatial resolution was sufficient to selectively measure light responses in the SCN.

Both in nocturnal and diurnal species, a proportion of the SCN neurons is inhibited rather than excited by light.^36^ In nocturnal species, the vast majority of neurons are light-excited, leading to excitatory responses at the population-level.^37^ In the diurnal degus and squirrel, however, the proportion of light-inhibited neurons exceeded the light-suppressed neurons.^38,39^ The function of light-inhibitions in the SCN has never been questioned, and has not been included in current models of photic entrainment. Given our finding that light causes an inhibitory response in the human SCN, we hypothesize that the functioning of the neuronal network is different between nocturnal and diurnal species. Mechanistic studies will be required in order to further dissect differences between diurnal and nocturnal species with respect to processing light information.

In summary, we report evidence that the circadian clock in humans receives multiple spectral inputs. This finding suggests that strategies designed to minimize the harmful effects of jet lag, rotating shift work, and other disruptions to our circadian rhythms should not exclusively focus on controlling blue light exposure, as other colors also affect the circadian clock. Moreover, our findings lead to a novel proposition that the clock of diurnal species is inhibited by light and thus that different neuronal circuitry within the SCN separates nocturnal and diurnal species. Our increasing knowledge regarding the effects of light on the circadian clock will definitely help responsible use of light in order to improve our general health and well-being.

## Methods

### Participants

This study was approved by the Medical Ethics Committee at Leiden University Medical Center, and all participants provided written informed consent. To be included in the study, the participants: *i*) were not color-blind, *ii*) had never had cataract surgery, *iii*) were free from internal metallic fragments or implants, *iv*) had no history of photosensitivity or seizure activity, *v*) had no history of sleep disorders, and *vi*) did not travel between time zones within 7 days prior to the scan. All 12 participants were males 30±12 years of age (range: 22-58 years). The Munich Chronotype Questionnaire was used to determine the participant’s midpoint of sleep (MSFsc).^40^ All data were collected from March 2019 through March 2020.

### Procedure

Participants were scanned between 10:00 pm and 11:00 pm. During the scan, the participants were instructed to move as little as possible, gaze upward without fixating on a certain point, and allow their eyes be exposed to the light. Both heart rate and breathing were measured continuously throughout the scanning session.

### Light exposure

Light was presented via a custom-made light-emitting device (Figure 1A and Supplemental Figure S1), which was attached to the head coil above the participant’s eyes in order to cover the participant’s field of view (Figure 1B). This device was programmed to emit monochromatic violet (λ_max_ = 405 nm), blue (λ_max_ = 470 nm), green (λ_max_ = 515 nm), or orange (λ_max_ = 590 nm) light via 4 sets of LEDs (see Figure 1C). The device’s aluminum controller box serves as a Faraday cage and receives input from an optical link with the computer located in the MRI control room. This controller box contains a trigger function in order to synchronize the LED’s protocol with the MRI sequence. Finally, any buildup of voltage in the LED device was prevented by including transient voltage suppressors, decoupling capacitors, current-limiting resistors, and pull-up resistors, and by shielding and twisting all wires (Supplemental Figure S1). The LED device was safety-approved in a Failure Mode and Effects Analysis. Non-dark-adapted participants were presented with 36 20-sec pulses of monochromatic light (9 pulses of each color) delivered in pseudorandom order in a block design; between each pulse, the participants were presented with 30 sec of darkness. Each light pulse included a gradual increase in intensity to avoid inducing a startle response (Figure 1D). The light stimuli were synchronized to the MRI sequence.

### fMRI data acquisition

Functional and structural images were acquired using a 7T MR scanner with a 32-channel head coil (Philips, Eindhoven, the Netherlands). High-resolution T1-weighted scans were acquired using the following parameters: field-of-view = 246×246×174 mm, flip angle = 7°, number of slices = 249, slice thickness = 0.70 mm, inter-slice gap = 0 mm, in-plane voxel resolution = 0.64×0.64 mm, echo time (TE) = 2.1 ms, repetition time = 4.6 ms and acquisition time = 506 sec.

Functional images were acquired using a T2-weighted BOLD fMRI sequence, with single-shot EPI (echo-planar imaging) and a repetition time of 641 ms. This temporal resolution is essential for reducing physiological noise using ICA-AROMA (Independent Component Analysis - Automatic Removal of Motion Artifacts) as described previously.^40^ Further scanning parameters were as follows: field-of-view = 180×16.4×180 mm, flip angle = 20°, A/P phase encoding direction, SENSE factor 2.8, slice thickness = 1.5 mm with an inter-slice gap of 0.15 mm, in-plane voxel resolution = 1.25×1.25 mm, echo time (TE) = 26 ms and acquisition time = 30.5 min. The functional images consisted of 9 or 10 slices in the transverse plane in order to achieve optimal spatial and temporal resolution within the partial field of view. These slices were aimed at the SCN (based on immunocytochemical staining with anti-vasopressin), the visual cortex V1 region, and the CSF. The dimensions of the SCN in adults are 1.1 mm (anterior-posterior) x 1.1 mm (left-right) x 1.7 mm (superior-inferior),^23^ which are similar to the voxel size of the functional scans. An additional whole-brain T2-weighted image using the same resolution as the functional time series data (1.25×1.25×1.65 mm) was also acquired in order to improve automatic registration for the partial field of view images.

### Pre-processing

The fMRI data were pre-processed using FSL 6.0.3 (FMRIB Software Library, Oxford, UK).^41^ The T1-weighted scans were extracted, followed by two-step registration to align the fMRI data to the high-resolution anatomical T1-weighted image and to the MNI152 1-mm template brain.

For the analysis of the SCN, manual registration of the T2-weighted to the T1-weighted scans was performed using ITK-SNAP (http://www.itksnap.org/), focusing on the region of the optic chiasm. In the second step, the T1-weighted scans were manually registered to the MNI152 standard space, followed by non-linear registration (10-mm warp resolution) using FSL FNIRT with a mask at the optic chiasm area that excluded large blood vessels. The average image for all functional (i.e., T2-weighted) scans transformed to MNI152 standard space clearly included the optic nerves (see the yellow rectangles in Figure 1H), indicating that manual registration was successful at aligning the relevant structures across participants and therefore accurately localized the SCN in all 12 participants.

For the analysis of the visual cortex, we performed automatic registration using FSL. Functional data were first registered to the T2-weighted low-resolution scan using 3 degrees-of-freedom (DOF) rigid-body transformation, then registered to the high-resolution structural scan using boundary-based registration (BBR);^42^ finally, the data were registered to the MNI152 template using 12 DOF linear transformation, followed by non-linear transformation with 10-mm warp resolution. The mean plot of the functional scans with automatic FSL registration (FLIRT and FNIRT) shows correct alignment of the large brain structures; however, the SCN appears as a gray blur (see Figure 1G), arguing that automatic registration is insufficient for analyzing small hypothalamic nuclei such as the SCN.

For the general linear model (GLM) analysis, we used spatial smoothing with a Gaussian kernel of 2 mm. No spatial smoothing was performed for the waveform analysis, thus maintaining the original data. For motion correction, Elastix, an open-source registration toolbox, was used with Euler (rigid) transformation parameters.^43^ To reduce noise in the data, artifacts were removed using ICA-AROMA, which automatically removes noise from the data by regression, without requiring previously acquired training datasets.^44^ The de-noised data were high-pass filtered at 0.01 Hz.

### Statistical analysis

Voxel-based GLM analysis was performed for each participant using Feat as implemented in FSL. Regressors were defined for each color separately and for all colors combined by convoluting the light protocols using a double gamma hemodynamic response function. Higher-level analysis was performed using the FLAME-1 mixed-effects model. The output z-stat map was set at a threshold of >2.3 or <−2.3 in order to identify voxels that showed a significant response to light. For the SCN analysis, a contrast of −1.0 was used to allow for characterization of the negative BOLD response. The region of interest was small, predefined and based on post-mortem anatomical data,^45^ we used a voxel-based analysis that was not corrected for multiple testing. The uncorrected data was used for illustration purposes (Figure 3, 4 A-D).

The analysis was spatially restricted to data extracted from an explicit mask. These SCN masks were drawn in standard space, based on an independent post-mortem brain dataset (preventing circularity). The regions of interest were thus defined *a priori*. For the SCN, bilateral masks (3×4×3 mm in size) were drawn using the MNI152 coordinates (mm): *x*: 84-86 and 93-95, *y*: 127-130, and *z*: 55-57. bilateral masks were also drawn for the CSF using the following coordinates (mm): *x*: 75-77 and 102-104, *y*: 125-128, and *z*: 52-54. Finally, bilateral masks were drawn for the visual cortex V1 region using the following coordinates: *x*: 85-87 and 91-93, *y*: 46-49, and *z*: 72-74. For each participant, the masks were transformed to native space, thresholded at 0.5, and binarized. Time series data from the transformed masks were extracted using the fslmeants command. Average waveforms were calculated by averaging the signal measured for each stimulus, synchronized to time 0 (i.e., the onset of the light stimulus). Data were normalized by dividing each data point in the average light stimulus by the mean baseline value (measured 15 seconds prior to the onset of the light stimulus) and then calculated as the percent change in the BOLD signal. To quantify the light response, we used the average percent change in the BOLD signal measured during the last 15 seconds of the light stimulus.

**Supplementary Information** is available for this paper.

## Acknowledgments

We thank Anouk Delhaas, Ilse Nootenboom, Huybert van de Stadt, Curtis Barrett, Thijs van Harten, Anne Hafkemeijer, Michel Villerius, Xu Chen, Jelle Goeman, Wouter Weeda, and Andrew Webb for technical assistance, help with data analysis, photography, editorial assistance and discussions. Post-mortem materials were obtained from the Netherlands Brain Bank. This research was supported by an ERC Advanced Grant (project number 834513 to J.H.M.) and the Velux Stiftung foundation, Zürich, Switzerland (project number 1131 to J.H.M.).

## Author contributions

R.A.S., L.H., C.R.W., L.S., M.J.P.v.O. and J.H.M. were responsible for the experimental design, J.A.M.J., S.v.B. and J.H.M. developed the LED device, D.F.S. performance and analysis of human histology, R.A.S., M.d.R., L.H., C.R.W., L.S., I.K., M.J.P.v.O. and J.H.M. acquired the fMRI data, R.A.S., M.d.R., L.H., C.R.W., L.S., I.K., M.J.P.v.O. and D.F.S. performed data analysis and R.A.S., M.d.R., J.A.M.J. and J.H.M. wrote the paper.

## Author information

The authors declare no competing interests. Correspondence should be addressed to J.H.M.

