## Supplemental information for "The photobiology of the human circadian clock"

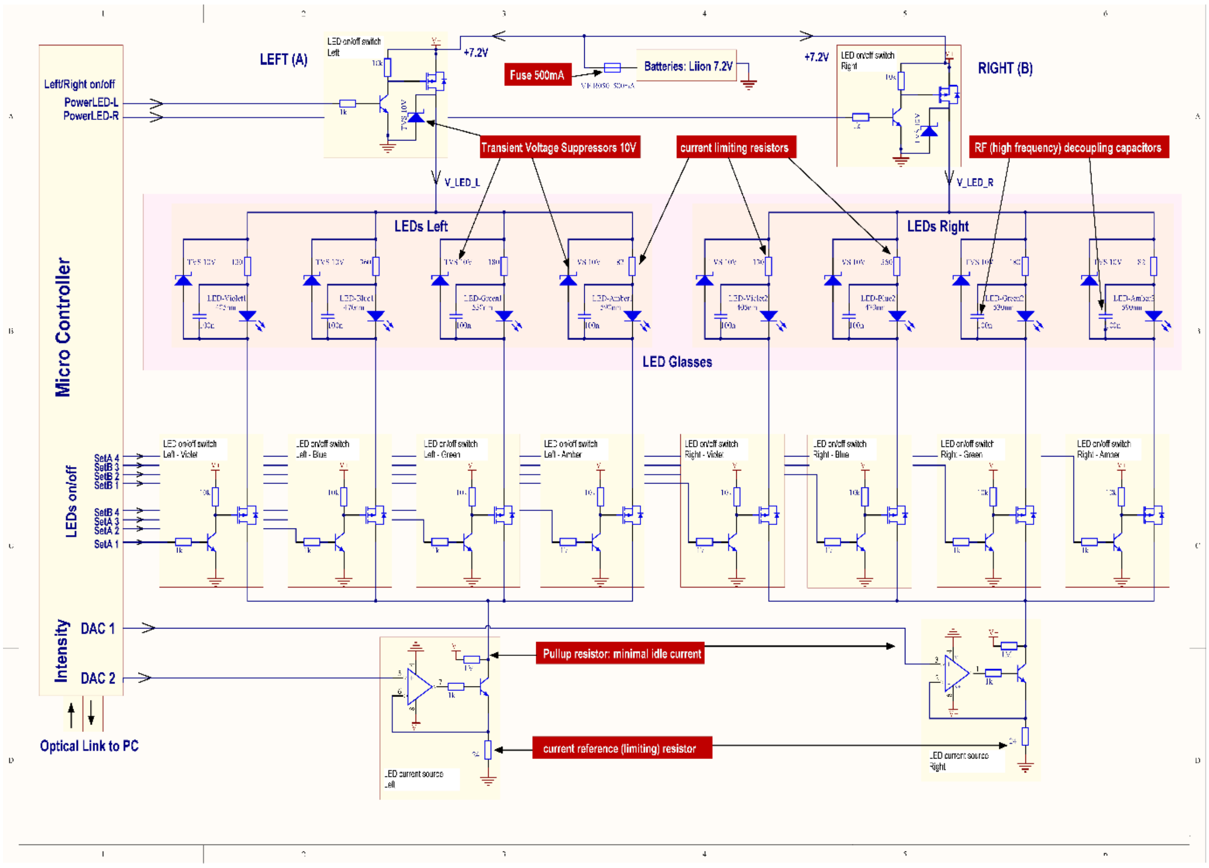


**Supplemental Figure S1: Circuit diagram depicting the MRI-compatible LED device.** The LED device was designed to be compatible with the high-frequency radio pulses and high magnetic field produced inside an MRI scanner. Moreover, we limited the use of ferromagnetic materials and used several strategies to prevent the buildup of voltage by the radio pulses (see text for details).

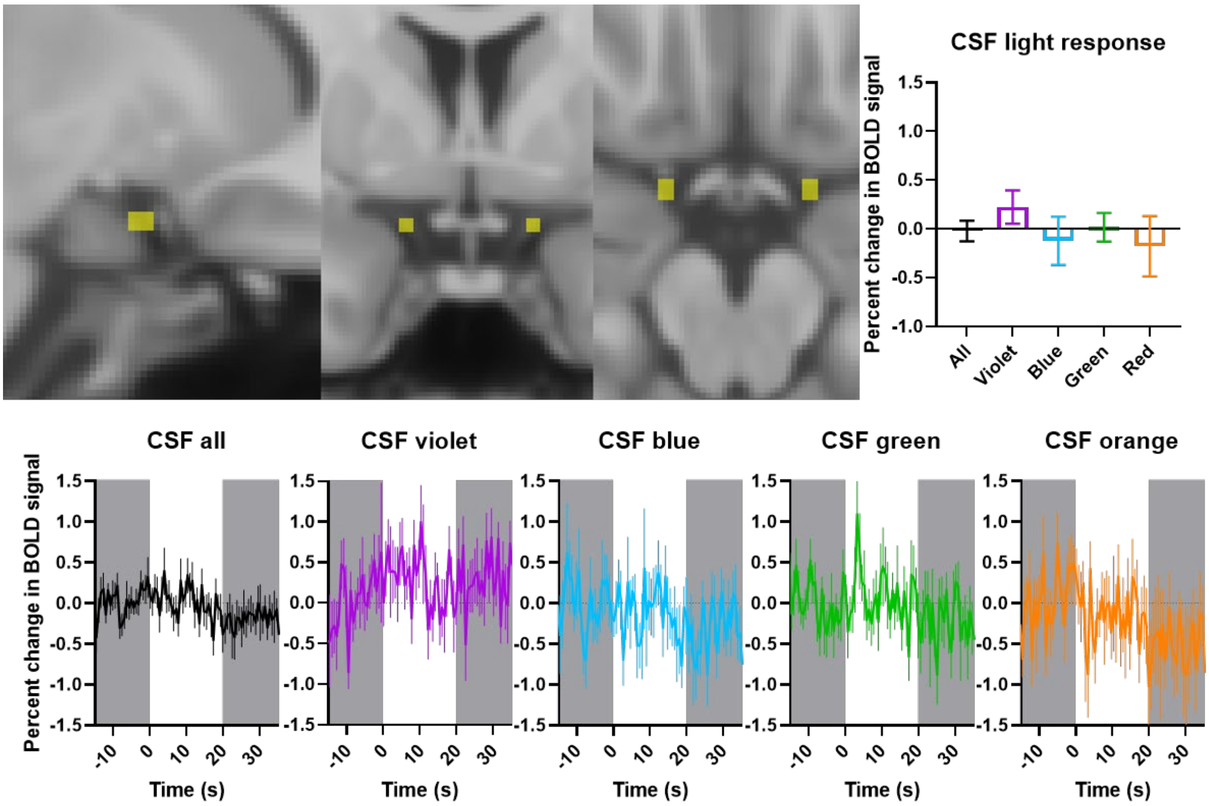


**Supplemental Figure S2: Light does not induce a change in the BOLD signal measured in the cerebrospinal fluid (CSF).** As a negative control, we analyzed the change in the BOLD signal measured in CSF surrounding the optic chiasm in response to violet, blue, green, or orange light (9 pulses each); "all" represents the average data obtained from all 36 pulses of light. The following coordinates (mm) of the mask were used: *x*: 75-77 and 102-104, *y*: 125-128, and *z*: 52-54.
